# Identifying Treatment Related Signatures In Glioblastoma Using KaleidoCell

**DOI:** 10.64898/2026.05.13.724848

**Authors:** Jean Radig, Carla Welz, Maria Jerome, Phil Sidney Ostheimer, Sophie Fellenz, Bernhard Radlwimmer, Carl Herrmann

## Abstract

Understanding how transcriptional heterogeneity is organized across tumors, patients, and treatment conditions remains a central challenge in cancer biology. Here, we present kaleidoCell, a GPU-accelerated Python framework for consensus non-negative matrix factorization that identifies reproducible meta-programs across independent samples. When benchmarked against its principal counterpart, the geneNMF R package, kaleidoCell achieves a twofold speed improvement on large datasets. In addition, it includes an integrated analysis module that generates a comprehensive HTML report containing key results and visualizations—including marker genes corresponding to the meta-programs, gene set enrichment analysis, UMAP projections and violin plots—without requiring additional user code. Using glioblastoma as a case study, we applied kaleidoCell to two published datasets. In a panobinostat-treated cohort, kaleidoCell resolves the cellular landscape of the tumor microenvironment and delineates how HDAC inhibition reshapes malignant cell states at single-cell resolution. We extend prior descriptions of the metallothionein-associated stress program in treatment response and identify co induction of IER3 as a candidate component of the associated survival signalling. In addition, we uncover novel transcriptional signatures associated with HDAC inhibition. Beyond confirming suppression of a neural progenitor cell-/oligodendrocyte progenitor cell-like program which is consistent with prior reports, kaleidoCell identifies loss of an astrocyte-like identity program as a previously unrecognized candidate mechanism of panobinostat action in glioblastoma. Together, these results establish kaleidoCell as a fast, user-friendly framework that enables robust discovery of biologically meaningful transcriptional programs in large, heterogeneous single-cell datasets.

## Introduction

Single-cell RNA sequencing (scRNA-seq) has transformed our ability to resolve cellular heterogeneity at unprecedented resolution. However, the high dimensionality and sparsity of the resulting data necessitate dedicated computational frameworks for meaningful interpretation (Luecken and Theis 2019; Hwang, Lee, and Bang 2018). A central challenge is the identification of latent transcriptional programs, reflecting underlying biological processes such as cell identity, cell state, or responses to environmental cues, from noisy, high-dimensional count matrices (Kotliar et al. 2019). Classical dimensionality reduction methods such as principal component analysis and independent component analysis have long been applied to this problem. However, the signed and typically non-sparse components they produce are often difficult to interpret biologically (Lee and Seung 1999). To address these limitations, several alternative approaches have been proposed. For example, single-cell hierarchical Poisson factorization (scHPF) Levitin et al. (2019) introduced a Bayesian framework that models transcript dropout and variable sparsity without requiring prior normalization. A consensus extension was later developed to improve robustness across random initializations (Levitin et al. 2023). Nevertheless, scHPF operates on a single expression matrix and does not provide a principled framework for identifying programs that are reproducible across independent samples. Other methods aim to improve interpretability by incorporating prior knowledge. Spectra (Kunes et al. 2024), for instance, encodes user-provided gene sets as a gene–gene knowledge graph and uses this structure as a regularization term to guide factorization. While this improves interpretability, it introduces dependence on the quality and completeness of the input knowledge base.

Non-negative matrix factorization (NMF) has emerged as a particularly powerful alternative for decomposing scRNA-seq data (Lee and Seung 1999; Lee and Seung 2001). By constraining both the basis and coefficient matrices to be non-negative, NMF yields a parts-based, additive decomposition that naturally produces sparse and interpretable gene programs. However, identifying biologically relevant transcriptional programs that are reproducible across samples remains a major challenge. Consensus NMF (cNMF) addresses this by performing multiple NMF runs at a fixed factorization rank on a single expression matrix and aggregating the results to derive robust gene programs (Kotliar et al. 2019). This approach has several limitations, including the high computational cost, the need to select a factorization rank k and the potential bias due to batch effects.

To overcome these constraints, Barkley et al. (2022) and Gavish et al. (2023) proposed a conceptually different strategy: here, NMF is performed independently on each sample, and the resulting programs are subsequently clustered across samples to identify recurrent signatures. This per-sample approach naturally mitigates batch effects and preserves sample-specific signals. The geneNMF package (Yerly et al. 2024) later made this strategy accessible through a user-friendly R toolkit. However, existing implementations are limited to R, restricting integration with Python-based workflows. In addition, neither framework provides integrated downstream analysis, which in practice constitutes a significant portion of the overall analysis effort.

Building on these foundations, we developed kaleidoCell, a GPU-accelerated Python framework for consensus NMF across multiple single-cell datasets. KaleidoCell addresses key limitations of existing approaches through several methodological contributions. First, rather than pooling samples into a single matrix, it performs NMF independently on each sample across a range of factorization ranks k, thereby eliminating the need for batch correction while preserving sample-specific signals and enabling cross-sample integration (Kotliar et al. 2019; Yerly et al. 2024). Second, in contrast to geneNMF, kaleidoCell implements a fully automated consensus strategy to derive the optimal number of meta-programs: candidate programs from all samples and ranks are compared using cosine similarity, clustered via Ward hierarchical clustering (Ward 1963), and evaluated using silhouette scoring (Rousseeuw 1987) to objectively determine the optimal number and quality of meta-programs, removing the need for manual parameter tuning. Third, the framework is implemented in Python with native GPU acceleration via PyTorch (Paszke et al. 2019), enabling efficient analysis of large datasets. Fourth, kaleidoCell streamlines downstream analysis by incorporating principled statistical and effect size–based filtering to identify meta-programs with meaningful differential activity across conditions (Squair et al. 2021). Finally, it provides a comprehensive suite of visualization tools, compiled into an intuitive HTML report, allowing researchers to efficiently explore and interpret the biological relevance of each identified meta-program.

## Results

### Kaleido Cell identifies robust transcriptional programs across samples

To extract robust and biologically meaningful transcriptional programs, we developed kaleidoCell, which integrates multiple NMF decompositions into unified meta-programs (Figure 1, Figure 2). The user provides a single-cell dataset along with a definition of what constitutes a sample for the decomposition step (Figure 2, step 1). This definition is flexible and context-dependent: for example, in a multi-patient study, samples naturally correspond to individual patients, enabling kaleidoCell to identify transcriptional programs that are reproducible across individuals.

**Figure 1:**
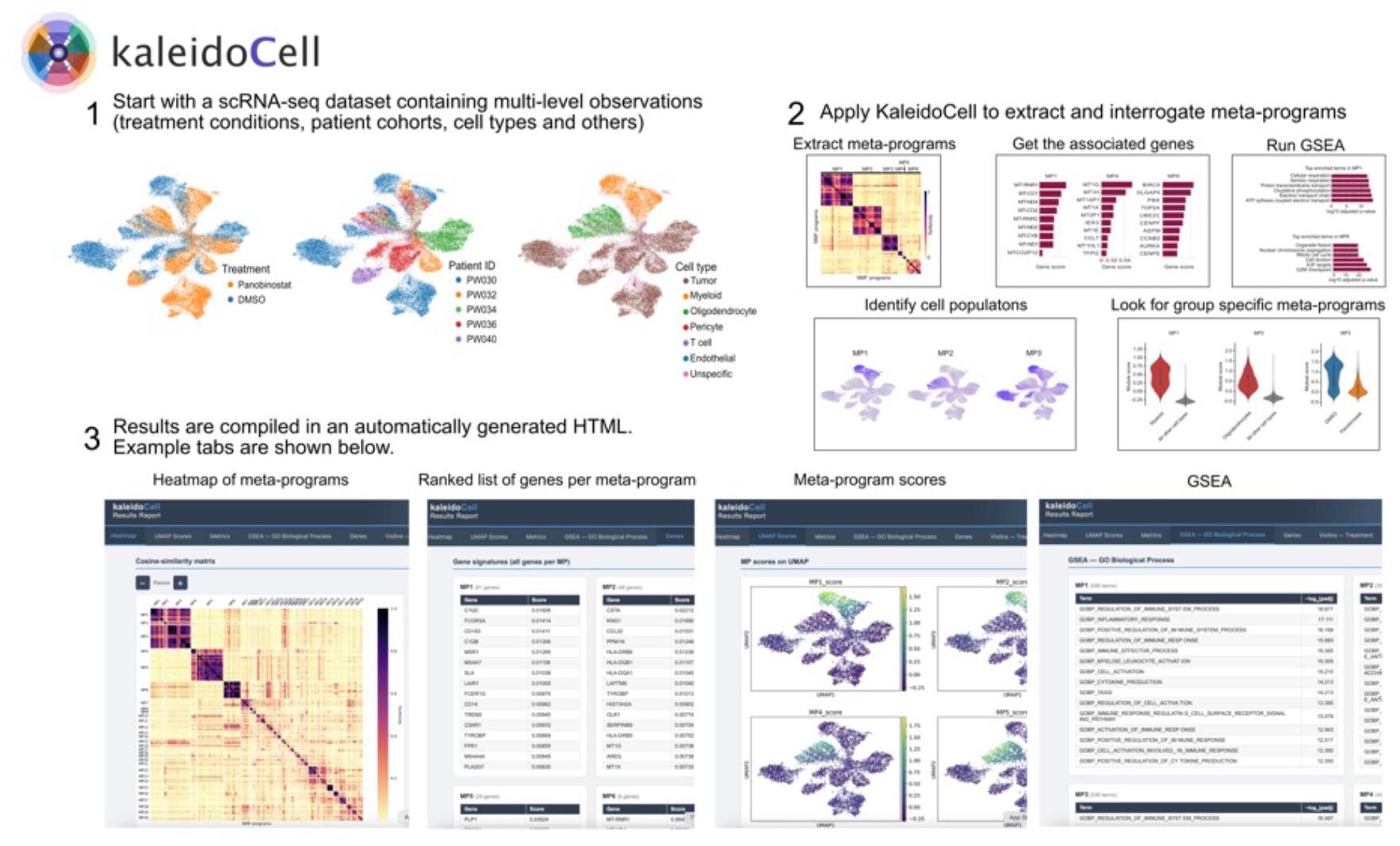
Overview of the kaleidoCell workflow. 1. The user provides a single-cell RNA sequencing dataset, potentially comprising multiple levels of annotation (e.g. treatments, patients or cell types). 2. kaleidoCell performs consensus analysis to identify meta-programs (MPs). The resulting MPs can be explored through an integrated analysis suite: users obtain the gene composition of each MP, associated GSEA results, and multiple visualization modalities. These include projection of MP activity scores onto UMAP embeddings and violin plots illustrating MP activity across user-defined groups, enabling intuitive characterization of cell populations. (3) All the results are aggregated in an HTML to ease subsequent reference and analysis.

**Figure 2:**
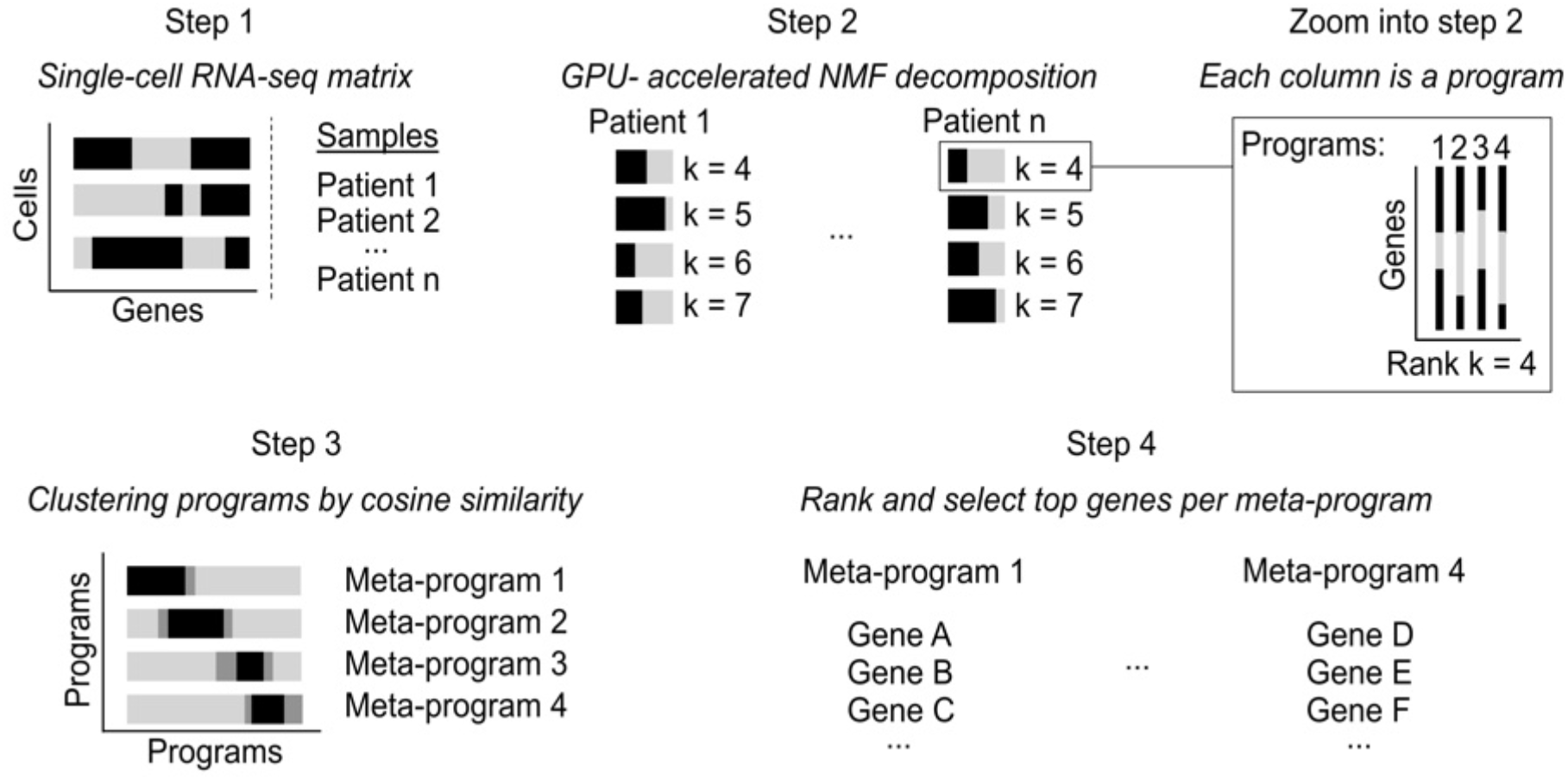
KaleidoCell’s main computational steps. Step 1. The user provides a single-cell dataset comprising multiple samples (e.g., patients) from which biologically meaningful transcriptional programs are to be extracted. Step 2. Each sample is independently decomposed using NMF to identify transcriptional programs. Zoom-in: each column of the resulting factor matrices corresponds to a transcriptional program; programs from all samples and factorization ranks are collected for downstream comparison. Step 3. Programs are compared across samples using cosine similarity to construct a program similarity matrix, which is then used to cluster related transcriptional signatures. Step 4. Within each cluster, gene weights are aggregated to compute consensus scores, and top-ranked genes are selected to define meta-programs representing shared transcriptional states across samples.

For each sample, NMF is run across a range of factorization ranks k (Figure 2, step 2). All resulting programs across all ranks are retained (Figure 2, zoom into step 2). Hence, kaleidoCell explores coarsegrained decompositions at low k and fine-grained decompositions at high k. Signatures that reflect genuine biological signals will tend to be recovered consistently across multiple ranks, while spurious or overfitted ones will not.

To identify reproducible patterns, all signatures are grouped by hierarchical clustering based on cosine similarity, effectively aggregating signals that recur across samples and factorization runs. Each resulting cluster defines a meta-program (MP) (Figure 2, step 3). Gene weights from each individual signature are averaged within each cluster, reducing noise from individual decompositions and emphasizing consistently co-expressed genes (Figure 2, step 4).

### A built-in reporting suite turns meta-programs into biological insight

A key limitation of existing tools is the lack of integrated, user-friendly downstream analysis. kaleidoCell addresses this by providing a self-contained visualization and interpretation suite that guides the user from raw MPs to biological insight (Figure 1).

Once MPs have been computed, kaleidoCell automatically generates an HTML report that can be browsed locally. The report is organized into several tabs that facilitate exploration of the results. A heatmap tab displays pairwise similarities between candidate programs, allowing assessment of cluster structure and potential manual refinement. A UMAP tab projects MP activity scores onto the cell embedding, enabling intuitive visualization of program activity across cell populations. A metrics tab summarizes per–MP quality indicators, including silhouette score, mean pairwise cosine similarity, and sample coverage (see Methods). A gene tab lists the top-ranked genes for each MP along with their consensus weights. A violin plot tab shows the distribution of MP activity scores stratified by user-defined observations (e.g., treatment condition or patient identity). A dedicated tab also provides gene set enrichment analysis (GSEA) results. Based on this report, users can iteratively refine the results by applying filtering criteria directly to the kaleidoCell object and regenerating the report on the filtered MPs.

### KaleidoCell retrieves biologically significant signatures in GBmap

To evaluate kaleidoCell on a large, well-characterised dataset, we applied it to a downsampled version of GBmap (Ruiz-Moreno et al. 2025), comprising 74,865 cells from 52 patients with cell-type annotations available at multiple hierarchical levels (Figure 3A,B; Supplementary Figure S5). Individual patients were used as separate samples for the kaleidoCell decomposition.

**Figure 3:**
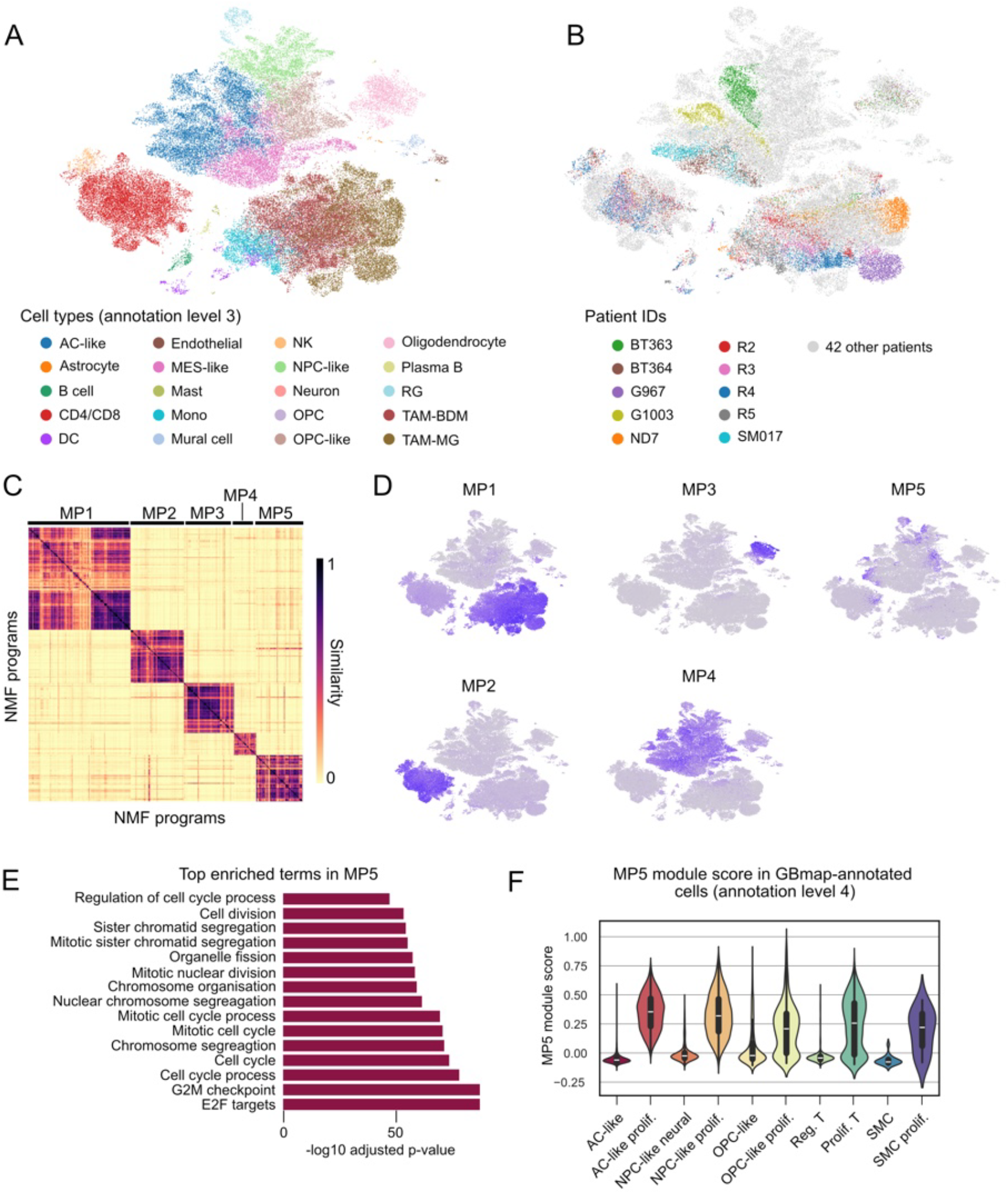
Application of kaleidoCell to a glioblastoma dataset (GBmap). A. Overview of GBmap annotations at the third level; annotations for levels one through four are provided in the Supplementary Figure S5. B. Patient composition in GBmap: the ten patients with the highest cell counts are shown, while the remaining 42 patients are not colored for clarity. C. Heatmap of meta-programs identified by kaleidoCell, after hierarchical clustering and selection of 5 meta-programs (MPs). D. Module scores for genes constituting the five MPs. E. GSEA on MP5, highlighting the top 15 enriched gene sets. F. Distribution of MP5 module scores across selected GBmap cells at annotation level 4.

Convergence and robustness of the results are checked, testing for different numbers of iterations to evaluate convergence, and iterations to evaluate stability. The results indicate low sensitivity to these parameters and fast convergence (Supplementary Figure S3, Supplementary Figure S4).

KaleidoCell identified thirty MPs using silhouette scoring. To focus on the most robust signals, we retained programs with high internal concordance and strong separation from other programs. We additionally required MPs to be supported by NMF programs originating from more than 30% of the patients in the dataset. Those filtering yielded a final set of five MPs (Figure 3C). Four of these (MP1-MP4) mapped cleanly to the major cell types present in the dataset (Figure 3D; Supplementary Figure S2), confirming that kaleidoCell reliably recovers dominant cell-type identity programs. The fifth MP (MP5) was active across multiple cell types. GSEA revealed strong enrichment for E2F targets, G2/M checkpoint, and cell cycle–related gene sets (Figure 3E). Consistent with this, MP5 activity scores were strongly associated with proliferating cells, as defined by GBmap level-4 annotations (Figure 3F; Supplementary Figure S5).

### KaleidoCell shows increased speed and meta-program definition compared to geneNMF on GBmap

Having established that kaleidoCell identifies biologically meaningful MPs across datasets, we next compared its performance to geneNMF, the method that inspired this work. We first evaluated computational efficiency. Using the GBmap dataset, we generated four subsets of increasing size by progressively subsampling cells and patients, and measured the runtime required to compute MPs. For smaller datasets, both methods performed comparably. However, as dataset size increased, kaleidoCell exhibited substantially improved scalability, achieving up to a two-fold reduction in runtime on the largest dataset (74 patients, 245,566 cells) (Figure 4A). We then compared the MPs identified by each method (Figure 4B). We constrained both approaches to produce 10 MPs. As geneNMF and kaleidoCell are stochastic methods, there may not be a direct one-to-one correspondence between programs. We used the Hungarian matching algorithm based on gene overlap to establish pairwise correspondences (Kuhn 1955). We first compared program sizes and observed that kaleidoCell generally produces MPs with a larger number of genes (Figure 4B), although this alone does not indicate biological relevance. To assess similarity between matched programs, we quantified the proportion of genes in the smaller set that are present in the larger set (Figure 4C). Using this metric, 8 out of 10 matched MPs showed at least 50% overlap, indicating substantial agreement between the two methods. To further evaluate the biological coherence of these programs, we compared their module score distributions across cells (Figure 4D). Overall, matched MPs displayed similar activity patterns, supporting their correspondence. Notably, kaleidoCell-derived programs often exhibited more specific and less diffuse activation across the dataset. This was particularly evident for MPs associated with distinct cell types, where kaleidoCell showed sharper localization of signal within relevant clusters. Importantly, kaleidoCell identified a lymphoid-associated MP (MP7) that was not recovered by geneNMF, suggesting improved sensitivity to certain biological signals. These results suggest that kaleidoCell recovers similar and potentially more refined, transcriptional programs compared to geneNMF with lower computational costs.

**Figure 4:**
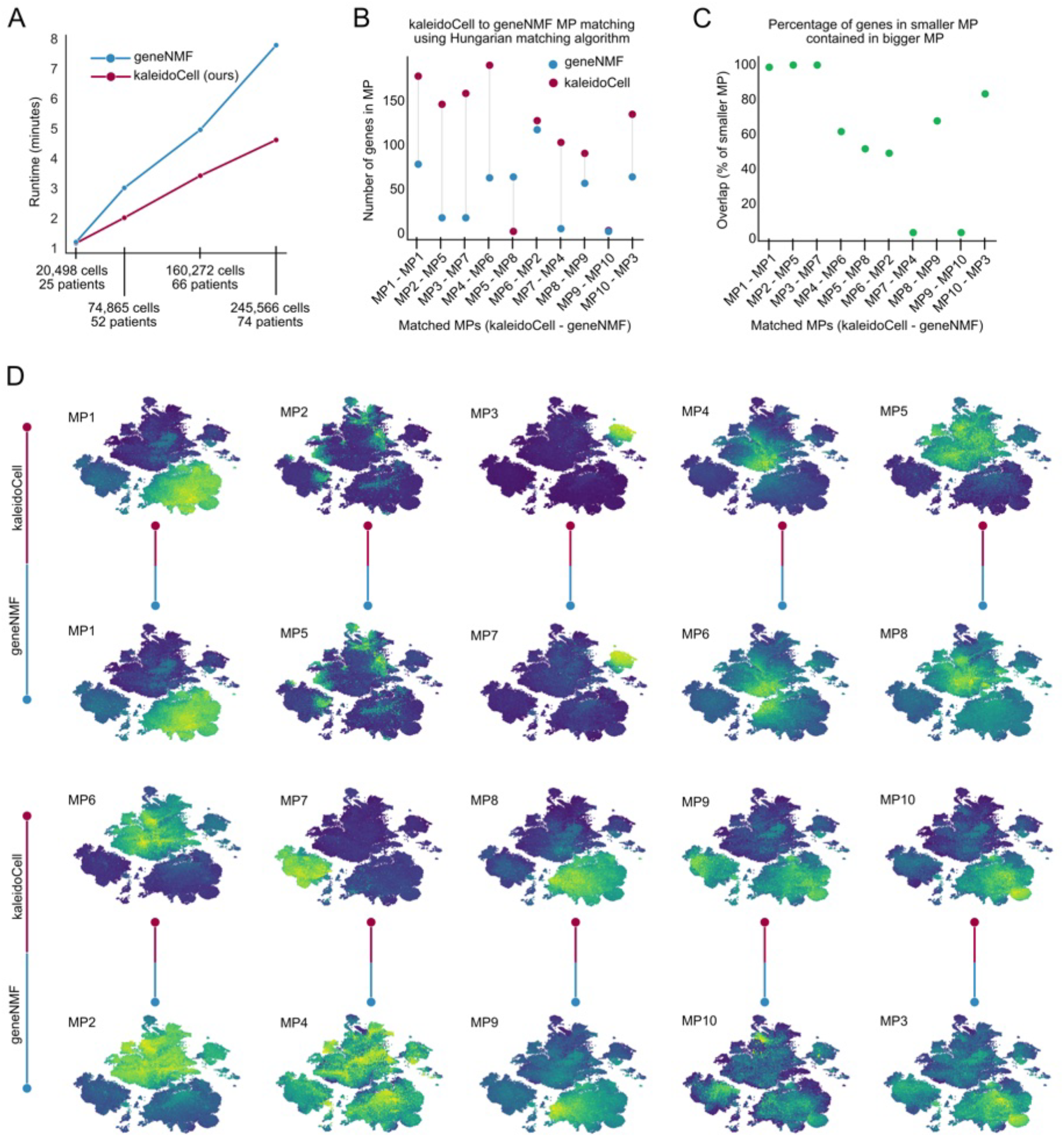
Comparative analysis of geneNMF and kaleidoCell meta-programs (MPs). A. Runtime comparison of geneNMF versus kaleidoCell across datasets with increasing numbers of cells and patients. B. Number of genes in kaleidoCell- and geneNMF-defined MPs, where 10 MPs were requested. MPs were matched using the Hungarian algorithm. C. Percentage overlap of genes between smaller and larger MPs, between matched MPs. D. Distribution of module scores for matched MPs, showing kaleidoCell scores aligned to the corresponding geneNMF MP.

### KaleidoCell rapidly decomposes a treated glioblastoma dataset into biologically interpretable meta-programs, recovering cell-type and treatment signatures

As an additional application, we evaluated kaleidoCell on a glioblastoma (GBM) single-cell dataset comprising both DMSO- and panobinostat-treated cells (Zhao et al. 2021; Levitin et al. 2023). The dataset includes samples from five patients and contains both malignant GBM cells and cells from the tumor microenvironment (Figure 5A–C), consistent with the complex cellular composition characteristic of GBM surgical specimens (Neftel et al. 2019; Patel et al. 2014). The dataset consists of 31,414 cells. Using kaleidoCell, we decomposed the dataset into a set of consensus transcriptional MPs in under one minute, highlighting the computational efficiency of the approach. For illustration, we focus on six representative MPs (Figure 5D,E). Two MPs reflect the transcriptional identity of non-malignant stromal cell types resident in the tumor microenvironment. MP1 is dominated by canonical myeloid and macrophage markers, including *TYROBP, C1QB/C, CD14, CD163, HLA-DRA/DRB1, TREM2, LYZ*, and *FCGR3A*, and represents tumor-associated macrophages and microglia, the dominant immune cell populations of the GBM microenvironment (Klemm et al. 2020; Müller et al. 2017) (Supplementary Table S1, Supplementary Material File 1, Figure S1A). GSEA confirmed enrichment for immune processes and inflammatory response, consistent with this identity (Figure 5F). MP2 captures the transcriptional program of mature oligodendrocytes and oligodendrocyte progenitor cells (OPCs), defined by *PLP1, MBP, MAG, MOG, CLDN11*, and *SOX10*, which are hallmark genes of myelin biosynthesis and oligodendrocyte differentiation (Marques et al. 2016) (Supplementary Table S1, Supplementary Material File 1, Supplementary Material File 1, Figure S1 B). These cells are non-malignant constituents of the brain parenchyma infiltrated by the tumor.

**Figure 5:**
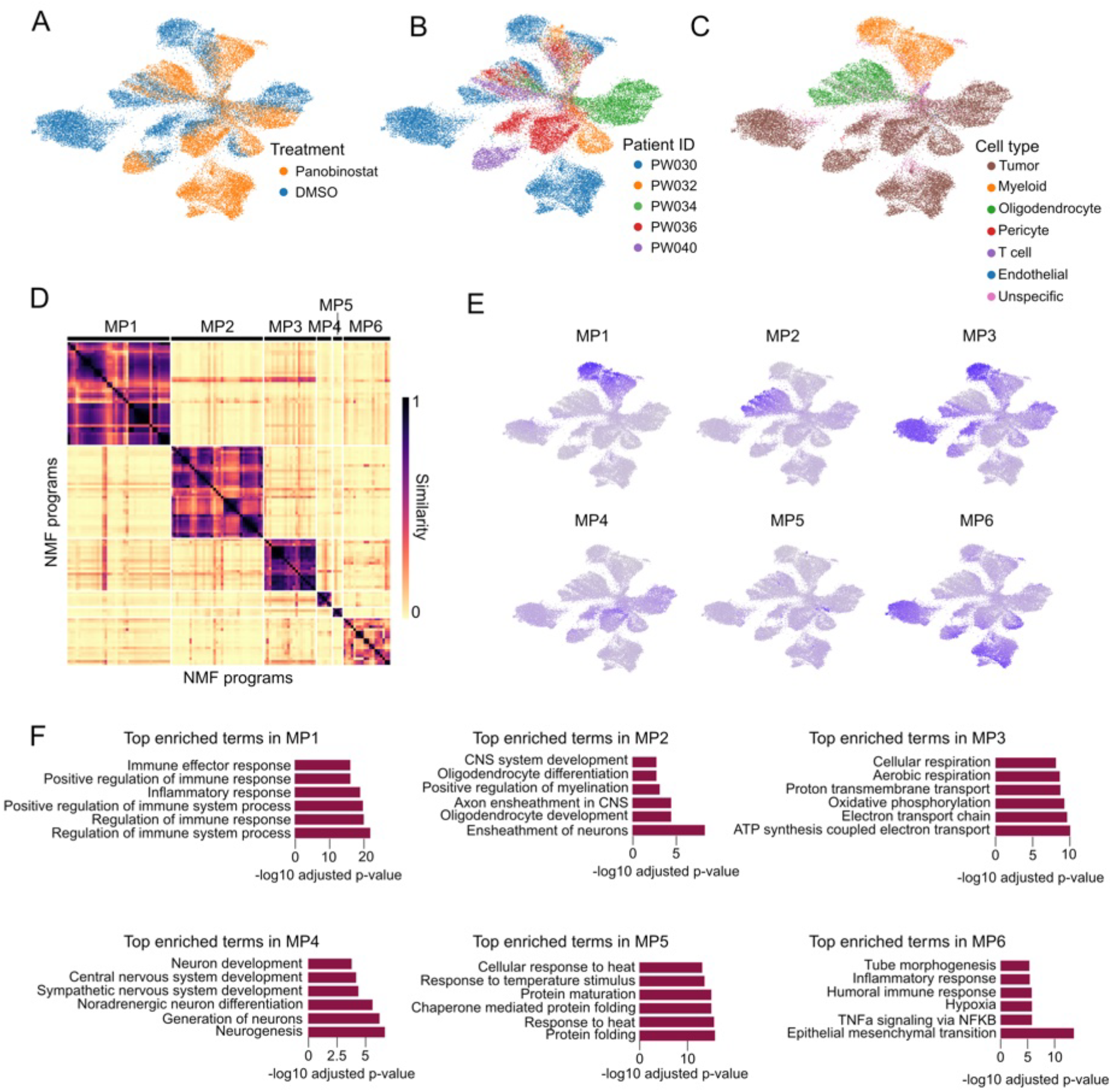
Application of kaleidoCell to a glioblastoma dataset (Peter Sims). A. UMAP representation of the dataset colored by treatment condition (DMSO vs. control). B. UMAP colored by patient of origin. C. UMAP showing annotated cell types. D. Heatmap of meta-programs (MPs) identified by kaleidoCell, after hierarchical clustering and selection of 6 MPs. E. UMAPs displaying MP scores. F. Gene set enrichment analysis of the six MPs, with the top six enriched terms shown.

The remaining MPs capture transcriptional states that are enriched in, but not exclusively restricted to, the malignant cell compartment. MP3 is dominated by mitochondrial transcripts (*MT-CO1, MT-ND1/2/4, MT-RNR1/2, MT-CYB*) and is associated with cellular respiration and oxidative phosphorylation (Supplementary Table S1, Supplementary Material File 1, Figure 5F). MP3 is detected across cell types including myeloid cells, consistent with a broadly shared metabolic state (Gavish et al. 2023) (Figure 5A, E). Moreover, MP3 activity is strongly suppressed upon panobinostat treatment, consistent with both the direct suppression of mitochondrial metabolic activity by panobinostat (Jane et al. 2023) and the loss of *MYC*-driven mitochondrial gene transcription upon HDAC inhibition (Li et al. 2005; Nebbioso et al. 2017; Barsotti et al. 2025) (Figure S1C).

MP4 is enriched for markers of neural progenitor cell (NPC) and OPC, including *SOX4, ASCL1, SOX11, DCX, DLL3, EGFR, OLIG1*, and *CCND2* (Supplementary Table S1, Supplementary Material File 1). This program closely mirrors the NPC- and OPC-like malignant cell states originally defined by Neftel *et al*., which represent a stem-like, proliferative axis of GBM heterogeneity (Neftel et al. 2019). MP5 is defined almost exclusively by inducible heat shock proteins and their co-chaperones, representing a stress response program. (Supplementary Table S1,Supplementary Material File 1). Finally, MP6 captures a broad inflammatory and extracellular matrix remodeling signature encompassing cytokines (*CXCL2, CXCL6, IL1B, LIF,CCL4/5*), acute-phase proteins (*SAA1/2, LCN2*), matrix remodeling factors (*FN1, TNC, COL6A1*), and NF-κB target genes (*ICAM1, BIRC3, NFKBIZ*) (Supplementary Table S1, Supplementary Material File 1). The concurrent upregulation of metallothionein stress genes (*MT1G, MT1X, MT2A*) further suggests oxidative stress as a contributing driver of this program. Together, the six MPs identified by kaleidoCell recapitulate the known cellular and transcriptional landscape of GBM, capturing both microenvironmental identities and tumor-intrinsic states.

### KaleidoCell recovers and extends known panobinostat signatures by resolving co-suppressed glioblastoma baseline programs

To investigate how the HDAC inhibitor panobinostat modulates transcriptional states in tumor cells, we restricted our analysis to the malignant cell compartment and directly compared DMSO (vehicle control) and panobinostat-treated conditions (Figure 6A).

**Figure 6:**
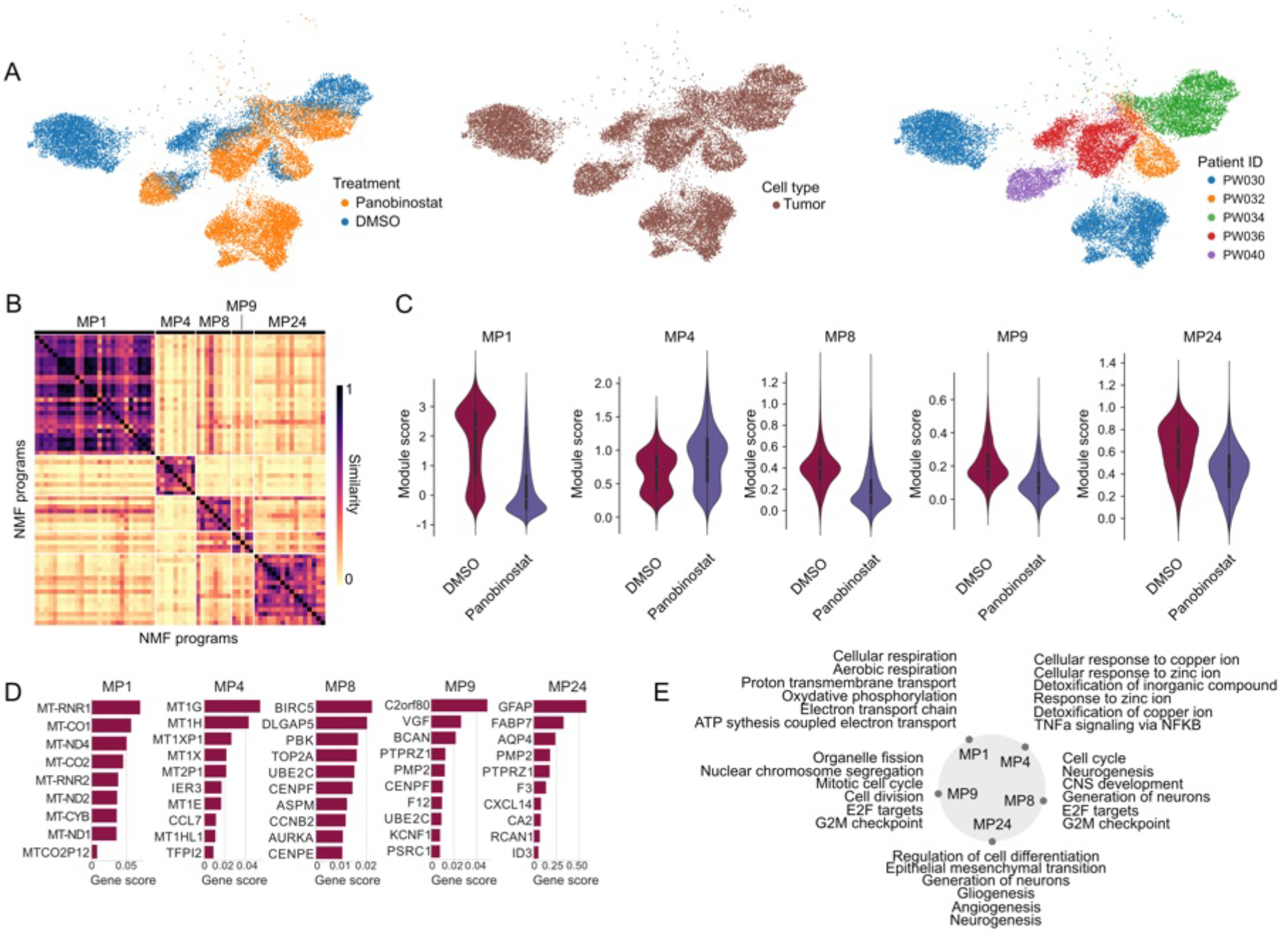
KaleidoCell analysis of tumor cells from the Peter Sims dataset. A. Subset of the dataset used as input for kaleidoCell analysis. B. Heatmap of the identified meta-programs (MPs) following filtering. C. Distribution of module scores for the retained MPs across cells. D. Top-ranked genes for each MP based on their weights. E. Top five enriched gene sets per MP.

Applied to the tumor-cell-only subset, kaleidoCell yielded thirty candidate MPs. To identify programs with meaningful differences in activity between conditions, we applied a two-stage filtering strategy. We applied an effect-size filter based on the Kolmogorov–Smirnov (KS) statistic, retaining only MPs with a KS value ≥ 0.20 to ensure a significant shift in activity. For the 13 programs passing this threshold, we applied three additional quality criteria. First, we required a silhouette score ≥ 0.20, reflecting a welldefined cluster boundary. Second, we required an average pairwise cosine similarity ≥ 0.30 among all NMF components contributing to a given MP, ensuring internal consistency. Third, to retain only signals reproducible across individuals rather than patient-specific artifacts, we required that each MP be represented by NMF components drawn from at least two of the five patients.

After this two-stage filtering, one MP (MP4) showed robust and consistent upregulation in panobinostat-treated tumor cells, while four MPs (MP1, MP8, MP9, MP24) were significantly enriched under the DMSO vehicle control condition (Figure 6B,C).

The panobinostat-enriched program, MP4, was dominated by metallothionein family members, which together accounted for approximately 53% of the total program weight (Table 1, Supplementary Material File 2, Figure 6D,E, Supplementary Figure S6). Beyond the metallothionein family members, our analysis revealed two co-induced genes absent from prior descriptions of this program, *IER3* and *CCL7. IER3*(immediate early response 3) is an NF-κB-regulated anti-apoptotic effector whose co-expression with metallothioneins at single-cell resolution suggests that panobinostat-treated cells may simultaneously engage metal-ion detoxification and survival signaling (Wu 2003). *CCL7* is also activated downstream of NF-κB, resulting in an inflammatory response and the recruitment of myeloid-dervied suppressor cells (Lou et al. 2021; Takacs et al. 2023; Song, Fu, Geng, et al. 2025).

**Table 1.**
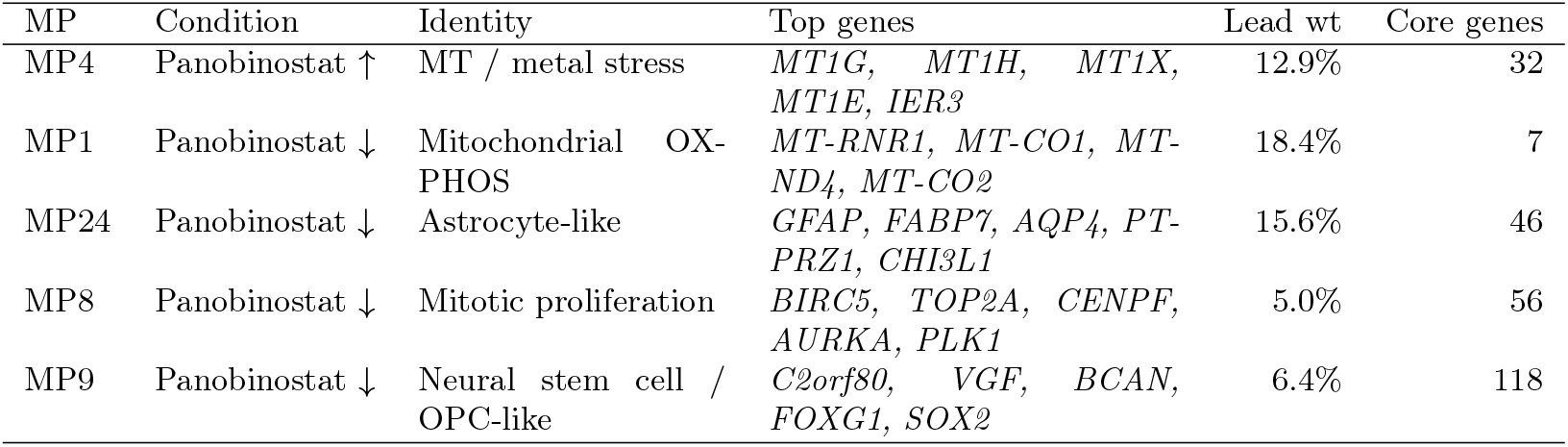
Summary of the five meta-programs identified by kaleidoCell in the DMSO∓panobinostat comparison. Lead weight: fraction of total program weight carried by the top-ranked gene. Core genes: number of genes required to cumulatively account for 80% of program weight.

The four panobinostat-repressed programs each correspond to a distinct aspect of the untreated GBM state (Table 1, Supplementary Material File 2, Figure 6D,E, Supplementary Figure S6). The high prevalence of mitochondrial genes in MP1 is similar to MP3 in the previous analysis (Jane et al. 2023; Li et al. 2005; Nebbioso et al. 2017; Barsotti et al. 2025). Enrichment for MP24 was dominated by canonical markers of the astrocyte-like (AC-like) transcriptional state in GBM as defined by Neftel et al. (2019) (Table 1, Supplementary Material File 2, Figure 6E, Supplementary Figure S6). GO enrichment for neurogenesis, gliogenesis, and glial cell differentiation confirms that MP24 represents a lineage-identity program rather than a generic stress response. MP8 captured a canonical mitotic proliferation program led by *BIRC5* (survivin, 5.0% of weight), *TOP2A, CENPF, AURKA*, and *PLK1*, with GO enrichment for mitotic cell cycle, chromosome segregation, and cell division (Table 1, Supplementary Material File 2, Figure 6D,E, Supplementary Figure S6).

Finally, MP9 reflected a heterogeneous NPC/OPC-like cycling state, with high-weight genes including *VGF, BCAN, FOXG1, SOX2*, and *OLIG1* co-occurring alongside cell-cycle markers such as *CENPF* and *TOP2A*, consistent with the cycling OPC-like subpopulation described in GBM single-cell hierarchies (Patel et al. 2014; Couturier et al. 2020).

## Discussion

In this work, we present kaleidoCell, a computational framework for extracting robust, consensus transcriptional programs from single-cell RNA-seq data through iterative NMF decomposition and hierarchical aggregation into meta-programs (MPs). Across two distinct GBM datasets, kaleidoCell consistently recovered biologically interpretable signatures that align with established cell-type identities and known transcriptional states, while offering meaningful advantages in computational efficiency and program resolution over existing approaches.

A central design principle of kaleidoCell is the exploitation of biological replication across samples. By running NMF independently per sample and across a range of factorization ranks, then clustering the resulting signatures by cosine similarity, kaleidoCell effectively distinguishes signals that recur reproducibly from those that are patient-specific or arise from overfitting. Our benchmarking on the large-scale GBmap dataset demonstrates that kaleidoCell achieves substantially faster runtimes as dataset size scales, with up to a two-fold reduction in computation time on datasets exceeding 200,000 cells compared to geneNMF. This scalability is important as single-cell atlases continue to grow in scope, and positions kaleidoCell as a practical tool for population-scale studies.

Beyond computational performance, the programs recovered by kaleidoCell show improved specificity relative to geneNMF. While the two methods agree on the majority of MPs, kaleidoCell programs exhibited sharper activity distributions, particularly for cell-type identity programs, and recovered a lymphoid-associated MP that geneNMF failed to detect.

The application to the panobinostat-treated GBM dataset illustrates the utility of kaleidoCell for dissecting drug-induced transcriptional remodeling at single-cell resolution. The single panobinostat-induced MP, dominated by metallothionein family members, recapitulates the signature previously reported in the same experimental system (Zhao et al. 2021; Levitin et al. 2023), validating our approach against an established benchmark. Beyond this replication, kaleidoCell identifies the co-induction of *IER3* alongside the metallothionein cluster. This co-induction is mechanistically interpretable: HDAC inhibitor treatment is a known activator of *NF-κB* signaling (Kim et al. 2006; Domingo-Domènech et al. 2008), and *IER3* is a direct transcriptional target of *NF-κB* whose protein product protects cells from apoptosis induced by *Fas* and *TNF-*ε (Wu 2003). *IER3* induction is therefore a by-product of the *NF-κB* defense response triggered by panobinostat itself. The net result is that individual panobinostat-treated GBM cells appear to simultaneously engage two distinct cytoprotective programs: metallothionein-mediated metal-ion detoxification on the one hand, and *NF-κB*-driven, *IER3*-mediated suppression of apoptosis on the other. The co-occurrence of these two responses at single-cell resolution raises the possibility that they act in concert to limit the cytotoxic efficacy of HDAC inhibition, and may partly underlie resistance to panobinostat.

The four panobinostat-repressed programs collectively map to distinct aspects of the baseline GBM transcriptional landscape: oxidative phosphorylation and mitochondrial respiration, an AC-like lineage identity program, a canonical mitotic proliferation program, and a cycling NPC/OPC-like state. Each corresponds to a well-characterized feature of the untreated tumor, and their coordinate suppression reflects the broad epigenetic reprogramming that distinguishes pan-HDAC inhibition from more targeted agents.

The repression of the AC-like program is consistent with recent single-cell evidence that AC-like glioma stem cells are especially sensitive to panobinostat, which induces a notable transition away from the AC-like state (Cirigliano et al. 2025). Downregulation of the mitotic proliferation program (MP8) is similarly expected: HDAC inhibitors suppress expression of G2 checkpoint kinases *WEE1* and *CHK1*, as well as key mitotic regulators including Survivin, causing glioma cells to enter mitosis with unrepaired damage and undergo mitotic catastrophe (Cornago et al. 2014). Finally, repression of the cycling NPC/OPC-like program (MP9) is consistent with the observations made by Levitin et al. and with the anti-stem-cell activity of HDAC inhibitors in GBM, as HDAC inhibitors exert cytotoxic and antiproliferative effects on sphere cultures enriched in glioma stem cells (Was et al. 2019), targeting precisely the progenitor-like, proliferatively active population proposed as a reservoir for tumor recurrence (Patel et al. 2014; Couturier et al. 2020; Cornago et al. 2014).

Several limitations warrant consideration. First, the NMF decomposition is inherently stochastic, and although kaleidoCell mitigates this through multi-run aggregation, the final set of MPs can be sensitive to the choice of filtering thresholds for silhouette score, cosine similarity, and sample coverage (Supplementary Figure S4). The built-in reporting suite enables iterative refinement, but the selection of appropriate thresholds will require biological judgment and may vary across datasets. Second, while kaleidoCell identifies transcriptional programs, it does not itself infer causal regulatory structure or assign directionality to relationships between programs. Integration with transcription factor activity inference or regulatory network methods would be a natural extension. Third, the current benchmarking focuses on GBM datasets; performance on tissues with different cellular complexity or lower between-sample reproducibility remains to be evaluated systematically.

Taken together, kaleidoCell provides a scalable, interpretable, and analytically self-contained framework for unsupervised discovery of transcriptional programs in heterogeneous single-cell datasets. Its combination of multi-rank, multi-sample NMF aggregation with an integrated downstream reporting suite addresses practical limitations of existing tools and offers a robust foundation for both exploratory atlas analyses and hypothesis-driven studies of transcriptional remodeling.

## Methods

### NMF decomposition of sample-level gene expression matrices

To decompose gene expression matrices into biologically interpretable transcriptional programs, we implemented a GPU-accelerated NMF framework in PyTorch (Paszke et al. 2019), using the multiplicative update scheme of Lee and Seung (Lee and Seung 2001). The NMF computation part is based on a reimplementation of the algorithm used in ShinyButchR to python (Quintero et al. 2020). NMF approximates a non-negative data matrix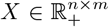, where n is the number of genes and m the number of cells, as the product of two low-rank non-negative matrices:

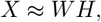

where 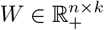 is the gene loading matrix, 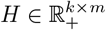 is the cell exposure matrix, and k is the factorization rank. The objective is to minimize the Frobenius norm of the reconstruction error:

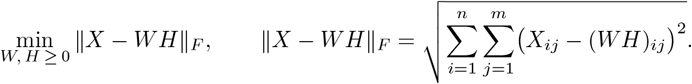

#### Optimization strategy and rank selection for NMF

1. Multiplicative update rules. NMF is solved iteratively via the multiplicative update scheme (Lee and Seung 2001): 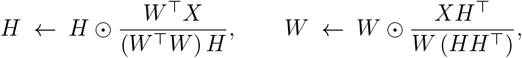

where ⊙ denotes element-wise multiplication and all divisions are performed element-wise.
2. Multiple initializations. To reduce sensitivity to local minima, n_init_ independent runs can be performed with random non-negative initializations of *W* and *H*. Each run proceeds until convergence, defined as a fixed number of consecutive iterations in which the assignment of each column of *H* to its dominant component does not change. Empirically, across both a GBM treatment dataset and GBmap, different initializations consistently converge to the same solution (Supplementary Figure S3B). Given the associated computational cost, the default is *n*_*init*_ = 1; users may increase this value if cluster instability is observed.
3. Convergence monitoring. At each iteration, the relative reconstruction error is computed as: 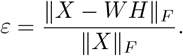 When multiple initializations are used, the factorization with the lowest ℰ is retained. The default number of iterations is 100; on the GBmap dataset, over 85% of runs reach at least 95% of their final convergence value by this point, indicating a favorable balance between computational cost and accuracy (Supplementary Figure S3A).
4. Multi-rank analysis. Factorization is performed over a range of ranks *k k*_1_, *k*_2_,. .., *k*_*r*_, reflecting different assumptions about the number of underlying transcriptional programs. Varying k modulates the granularity of the resulting meta-programs while the overall program structure remains largely stable (Supplementary Figure S4). We recommend exploring a range spanning three below to three above the anticipated number of cell states or clusters. By default, k ranges from 4 to 9.

### Clustering of transcriptional programs into consensus meta-programs

After computing NMF decompositions for each sample, consensus meta-programs are derived by clustering transcriptional programs across samples using pairwise similarity and extracting representative genes per cluster.

Let *S* denote the number of samples, n the number of genes, and 𝒦 = {*k*_*1*_,. .., *k*_*R*_} the set of factorization ranks applied to every sample (default *K* = {4, 5, 6, 7, 8, 9}). For each sample *s* ∈ {1,.. ., *S*} and each rank *k* ∈ 𝒦, NMF produces a gene loading matrix 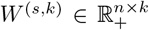, whose j-th column 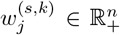 represents one transcriptional program (Figure 2 Step 2, zoom into step 2). The full set of programs for sample s is obtained by concatenating across ranks:

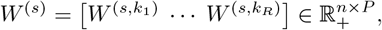

where 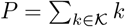 is the number of programs per sample.

Before concatenation across samples, each *W* ^*(s)*^ is transformed independently to suppress broadly expressed genes and amplify state-specific ones. For gene g and program *j* within sample *s*, the per-gene specificity score is:

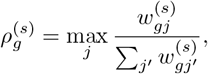

which is high when gene g dominates a single program and low when its weight is spread evenly. Each loading 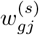 is rescaled by 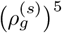 and the columns are 𝓁_1-_normalized, yielding the specificity-weighted matrix 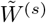 with columns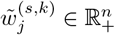.

All programs across all samples are then concatenated into a single matrix:

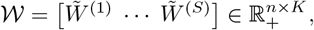

where *K* = *S·P* is the total number of programs. Each column of 𝒲 is one specificity-weighted transcriptional program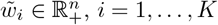.

Pairwise similarity between programs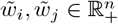 is measured by cosine similarity:

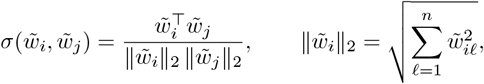

yielding a similarity matrix *C* ∈ [0, 1]^*K*×*K*^ with entries 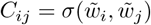.

1. Distance matrix. A distance matrix 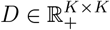 is derived from C by: 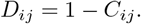
2. Finding the optimal number of meta-programs. Ward linkage hierarchical clustering (Ward 1963) is applied once to the distance matrix D, producing a full dendrogram. The dendrogram is then cut at each candidate value *K* ∈ {2,…, *K*_max_}(default *K*_max_ = 30), yielding a flat partition of all programs into K clusters. For each cut, the mean silhouette score (Rousseeuw 1987) is computed to assess cluster cohesion and separation, and the value K^*\**^ maximizing this score is retained as the optimal number of meta-programs. Users may override K^*\**^ and specify the number of clusters directly to incorporate prior biological knowledge.

### Extraction of representative marker genes per meta-program

Each cluster 𝒞κ groups a set of specificity-weighted programs 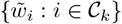, each being a vector of n gene weights. To compute the consensus score *ϕ*_*gk*_, one would naively average 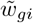 across programs. However, a single aberrant program could dominate the average for a given gene. To guard against this, an outlier masking step is applied per gene before averaging: for each gene g, the mean and standard deviation of its weights across the programs in the cluster are computed:

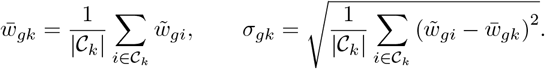

Any individual weight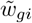 falling outside the interval 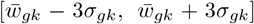 is masked and excluded from the average. This masking is only applied when |𝒞κ| ≥3, as fewer programs do not provide a reliable estimate of *σ*_*gk*_. The consensus score is then computed as the mean over non-masked values:

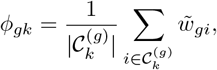

where 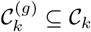 denotes the subset of programs not masked for gene g.

Kaleido Cell offers two methods for selecting marker genes from *ϕ*_*gk*_.

#### *KMeans-based selection* (default)

Genes are ranked by *ϕ*_*gk*_ and partitioned into two groups by k-means clustering (*k* = 2) applied to the score distribution. Genes assigned to the group with the higher mean score are retained as markers of the meta-program, avoiding the introduction of an arbitrary count threshold.

#### *Confidence-based selection* (employed by geneNMF)

Two sequential filters are applied. First, a weightcoverage threshold α ∈ (0, 1] retains only the top-ranked genes that collectively account for at least a fraction αε of the total consensus weight:

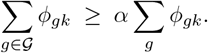

Second, a consistency threshold *β* ∈ (0, 1] requires that each retained gene ranks among the top genes in at least a fraction α of programs within the cluster:

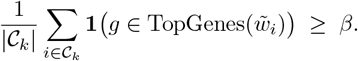

The gene set is capped at *G*_max_ = 200.

### Post-processing of the meta-programs

To ensure the biological relevance and robustness of the identified meta-programs, kaleidoCell provides a flexible post-processing framework combining quality-based filtering and statistical testing.

Low-quality meta-programs can be removed using intrinsic clustering metrics. Each meta-program is evaluated on three criteria. First, its mean silhouette score, computed by averaging the silhouette scores of all NMF programs assigned to the cluster, where each individual silhouette score measures how much closer that program is to the other programs in its own cluster than to programs in the nearest neighboring cluster, based on the pairwise distance matrix *D*. Higher values indicate a compact and well-separated cluster. Second, its mean intra-cluster cosine similarity, computed by averaging the pairwise cosine similarities between all NMF programs within the cluster, which reflects internal consistency in gene-weight profiles. Third, its sample coverage, defined as the fraction of biological samples contributing at least one NMF program to the cluster. Thresholds can be set for any combination of these metrics, with either permissive (logical AND) or stringent (logical OR) filtering. In this study, we required a mean silhouette score of at least 0.20 and a mean intra-cluster cosine similarity of at least 0.30.

Second, to identify condition-relevant meta-programs, kaleidocell evaluates differential activity across cell groups. Module scores are computed per cell and per MP using scanpy, and a two-sample Kolmogorov– Smirnov (KS) test is applied between the score distribution of the target condition and all remaining cells, with p-values corrected via the Benjamini–Hochberg procedure. Because large single-cell datasets can yield highly significant p-values even for negligible differences, we additionally require a minimum KS statistic D_KS_ ≥ 0.20, where D_KS_ is the maximum absolute difference between the two empirical cumulative distribution functions. Optionally, the Wasserstein distance can be used as an alternative effect-size metric. Directionality filters may also be applied to retain only MPs upregulated in the condition of interest.

### Datasets

#### Glioblastoma dataset with treatment

The GBM dataset with treatment comprises single-cell transcriptomic profiles of patient-derived GBM cells and is available under GEO accession number GSE148842. For our analysis, we used the annotated version provided by Levitin et al. (2023), accessible at https://drive.google.com/file/d/18-KInmm43wKdBX95Gq9xbuzAQwtLjgE9/view?usp=sharing. We subset the dataset to include five patients (PW030, PW032, PW034, PW036, and PW040) for which both control and panobinostat-treated conditions were available, excluding all other treatment groups. Specifically, this subset was defined by selecting the following entries from the ‘Sequencing Name’ column in ‘adata.obs’: [‘PW029-N701’, ‘PW030-N701’, ‘PW030-N702’, ‘PW030-N704’, ‘PW032-N702’, ‘PW032-N705’, ‘PW032-N709’, ‘PW032-N711’, ‘PW034-N701’, ‘PW034-N702’, ‘PW034-N705’, ‘PW036-N701’, ‘PW036-N702’, ‘PW036-N705’, ‘PW040-N701’, ‘PW040-N703’, ‘PW040-N705’, ‘PW040-N707’, ‘PW040-N709’, ‘PW040-N711’, ‘PW040-N712’].

When duplicated gene names were present, counts for the first occurrence were retained. Raw counts were normalized to 10,000 counts per cell and log-transformed using log_1_ p. The original dataset contains 81,054 cells (65,347 control and 15,707 perturbed) across 20,098 genes. To balance the comparison between conditions, control cells were randomly downsampled to match the number of perturbed cells. We further downsampled the number of genes to the top 7,000 most highly variable ones. The dataset used to run kaleidoCell on comprised 31,414 cells and 7,000 genes. This dataset can be found at https://doi.org/10.5281/zenodo.19918636 under name `gb.h5ad` (Radig et al. 2026).

#### GBmap dataset

The GBmap dataset was obtained from the CellxGene platform (Collection ID: `999f2a15-3d7e-440b-96ae-2c806799c08c`) (Ruiz-Moreno et al. 2025). The Core GBmap was used, provided as a preprocessed `h5ad` file with cell-type annotations available at four hierarchical levels (`annotation level 1` through `annotation level 4`). No additional preprocessing was performed, as GBmap is an already curated dataset; log_1_p-normalized expression values were used throughout, corresponding to the default layer. The dataset contains 338,564 cells and 27,983 genes and 110 patients. The patients are accessible through the observation donor id. The results presented in Figure 3 were computed on a subset of the dataset. We selected 25% of the cells, randomly sampled. We kept patients that had at least 500 cells. The dataset comprised 74,865 cells and 27,983 genes and 52 patients.

For the runtime benchmarking experiment, the dataset was subsampled at fractions of 0.10, 0.25, 0.50, and 0.75 of the total cell count using `numpy.random.choice`. In each subsampled dataset, donors represented by fewer than 500 cells were excluded to ensure suffcient per-sample coverage for factorization.

### Packages used for the computations

NMF decomposition was performed using PyTorch (Paszke et al. 2019), with data structures provided by AnnData (Virshup et al. 2023), NumPy (Harris et al. 2020), and pandas (McKinney 2010). Cosine similarities between NMF programs were computed with `sklearn.metrics.pairwise.cosine similarity` from scikit-learn (Pedregosa et al. 2011). Programs were hierarchically clustered using Ward linkage via `scipy.cluster.hierarchy.linkage` (method = `“ward”`) from SciPy (Virtanen et al. 2020), and cluster assignments were derived with `scipy.cluster.hierarchy.fcluster`. Meta-program quality was assessed using the silhouette score (`sklearn.metrics.silhouette samples`). Differential activity between conditions was tested with Kolmogorov–Smirnov test (`scipy.stats.ks 2samp`) or Wasserstein distance (`scipy.stats.wasserstein distance`), with multiple-testing correction applied using the Benjamini–Hochberg procedure via `statsmodels.stats.multitest.multipletests` (Seabold and Perktold 2010). GSEA was performed using gseapy (Fang, Liu, and Peltz 2023) via Enrichr (`gseapy.enrichr`). Using the MP gene scores, the user can also use the preranked GSEA (`gseapy.prerank`). Visualizations were produced with Matplotlib (Hunter 2007) and seaborn (Waskom 2021).

### Comparison against geneNMF

To benchmark kaleidoCell against geneNMF, we used the geneNMF implementation from GitHub at commit `7ac4752` (pushed 26 September 2025) with default settings. To ensure optimal GPU acceleration for both tools, all experiments were run within a dedicated Docker container, made available in the GitHub repository. Runtime was defined as the wall-clock time elapsed from the start of factorization to the extraction of MPs, excluding file loading and any post-processing steps.

To match meta-programs identified by geneNMF against those identified by kaleidoCell, we used the Hungarian matching algorithm (Kuhn 1955) as implemented in `scipy.optimize.linear sum assignment` (**Virtanen2020**). Let 𝒜 = {*A*_1_,. .., *A*_*m*_} and ℬ = {*B*_1_,. ..,*B*_*n*_} denote the sets of top genes defining each meta-program output by geneNMF and kaleidoCell, respectively. For each pair (*A*_*i*_, *B*_*j*_), we computed the overlap coefficient

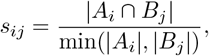

which quantifies the fraction of the smaller gene set covered by the intersection, and is thus robust to differences in meta-program size between the two tools. The resulting m n similarity matrix *S* = (*s*_*ij*_) was converted to a cost matrix *C* = **1** − *S*, and the optimal one-to-one assignment 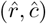 minimizing the total cost

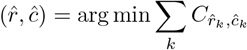

was obtained using the Hungarian algorithm, yielding the best global pairing between the meta-programs of the two methods. The same method was applied in Supplementary Figure S4 where we look at the robustness of kaleidoCell when varying the values of k.

## Code availability

The code is available on GitHub at https://github.com/JeanRadig/KaleidoCell. The package kaleidoCell can be installed using conda. We also provide a Dockerhub-available docker image for CUDA 12.4. The docker image can be fetched from hdsu/kaleidocell env:latest. We provide the source Docker file, such that the Docker can be adapted to any CUDA version.

## Author contribution

JR designed the study. JR, CW and PSO developed the program. JR and MJ conducted and evaluated the different experiments. CH provided guidance and feedback throughout the course of the project. All authors read and approved the final manuscript.

## Acknowledgments

JR, MJ and PSO are supported by the Carl-Zeiss-Stiftung through the AI-CARE project (P2022-08-008).

## Supplementary Figures

**Figure S1:**
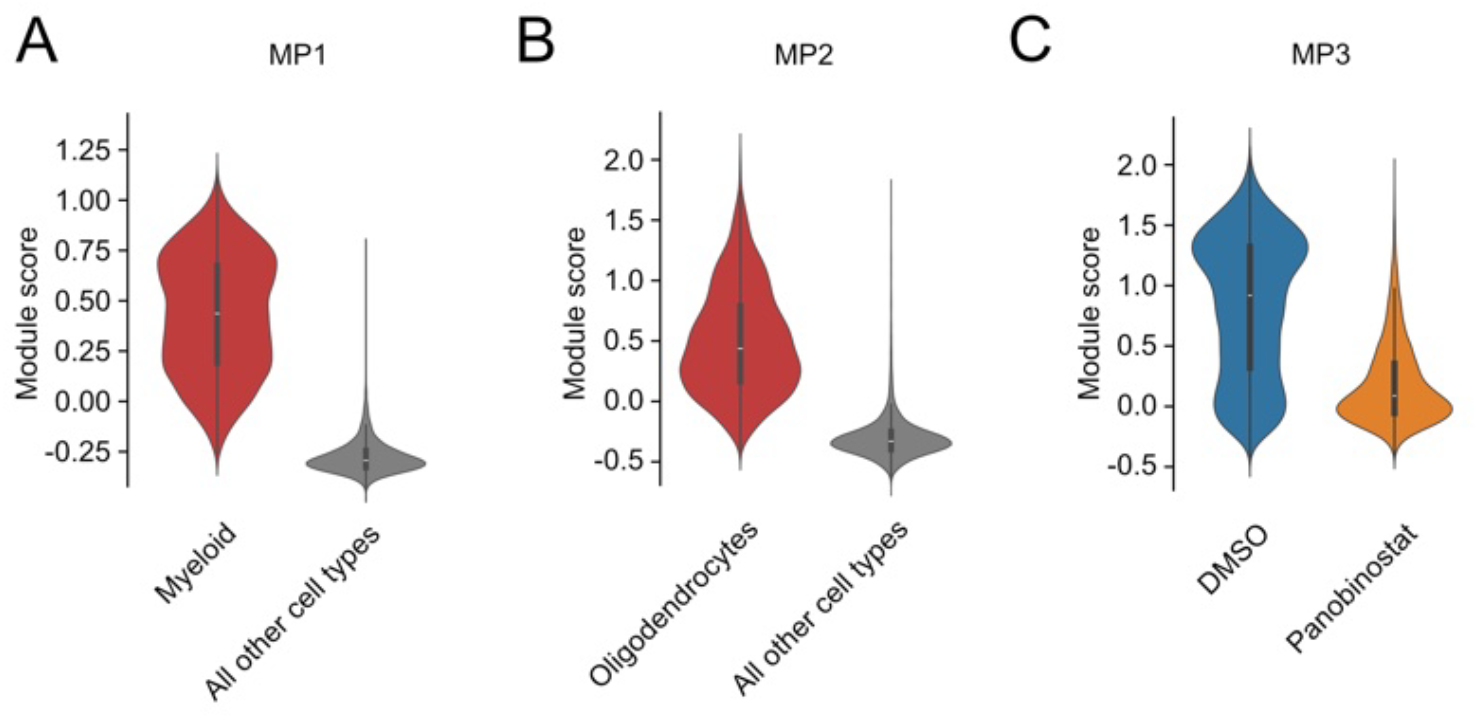
Activity distribution in the Peter Sims dataset. A. Distribution of MP1 module scores across cell types. B. Distribution of MP2 module scores across cell types. C. MP3 module score across treatment conditions. MP = meta-program.

**Table S1.**
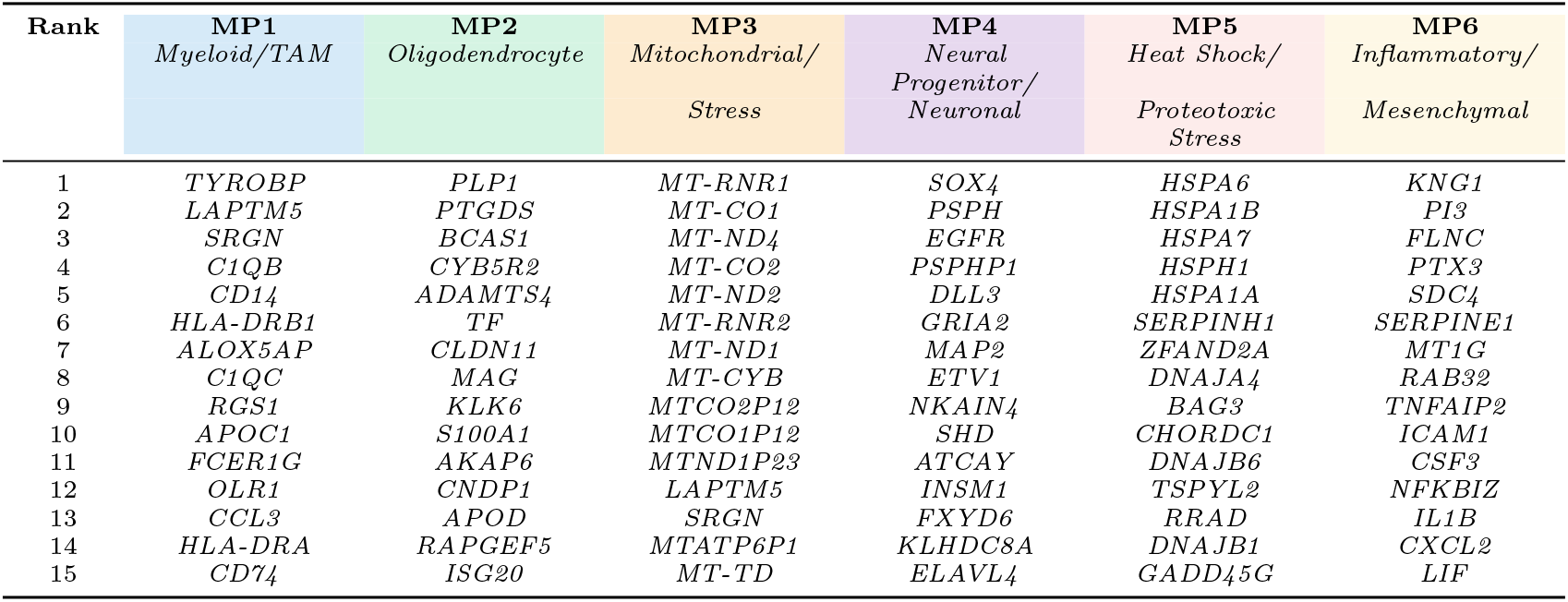
Top 15 genes for each consensus meta-program (MP) (MP1–MP6) identified by kaleidoCell on Peter Sims. Genes are ranked by their consensus weight within each meta-program, with rank 1 being the highest-weighted gene. MP identities were assigned based on GSEA enrichment and manual inspection of top-ranking genes.

**Figure S2:**
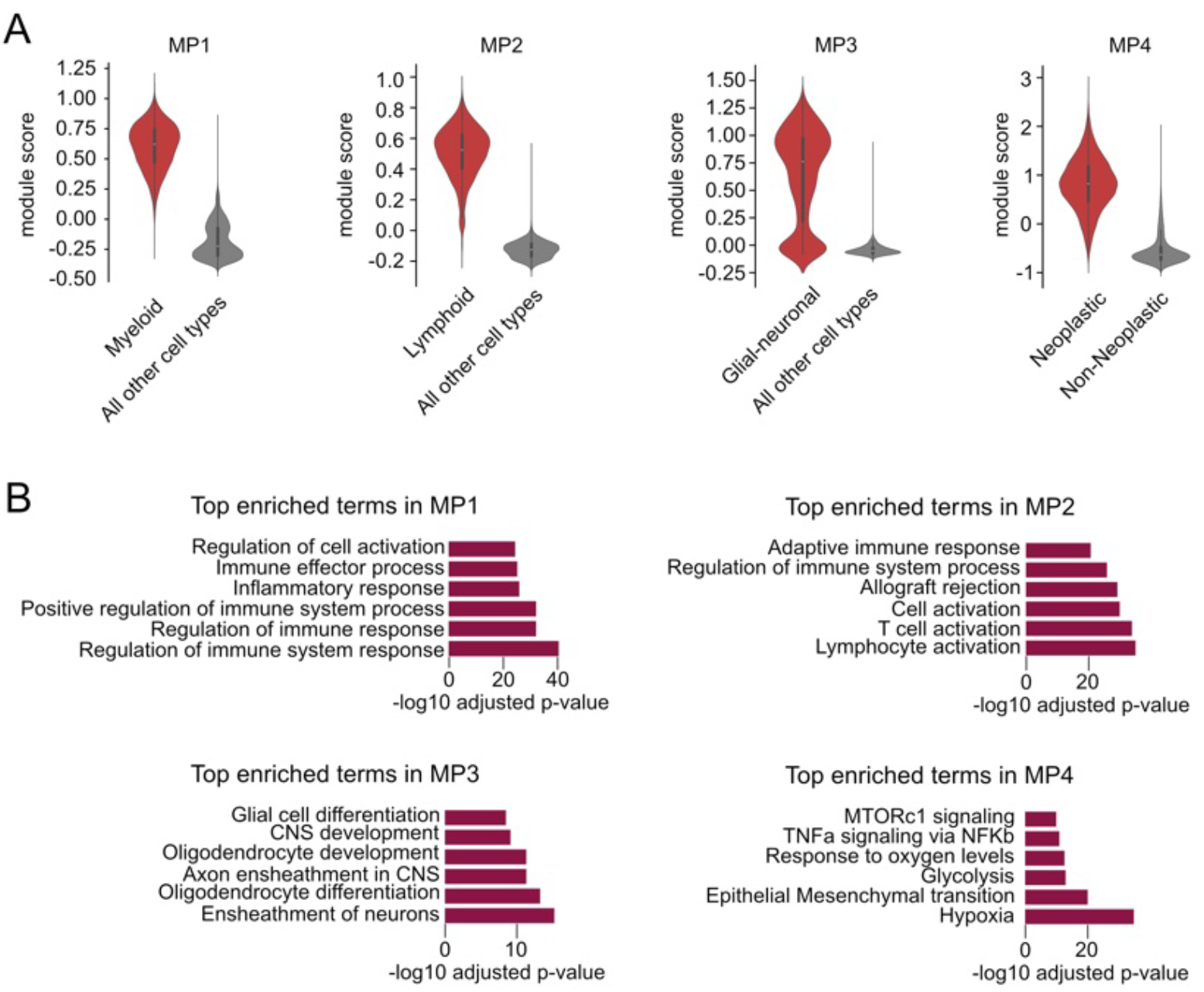
Characterization of meta-programs (MPs) from the GBmap dataset. A. Module scores for MPs 1–4, illustrating the relative enrichment of each meta-program across cell types. B. Gene set enrichment analysis for MPs 1–4, highlighting associated biological pathways and functions.

**Figure S3:**
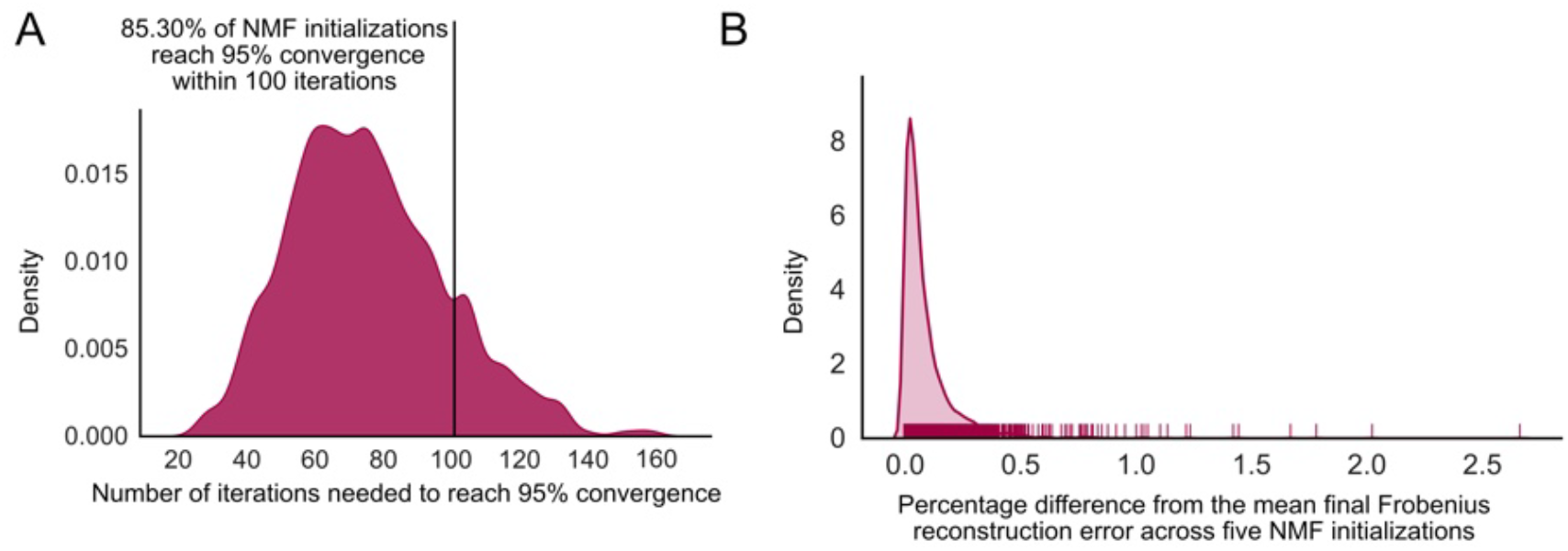
Convergence characteristics of kaleidoCell on GBmap. A. Number of iterations required to reach 95% convergence. B. Variation in final Frobenius reconstruction error across five independent NMF initializations. Each NMF run was performed five times, and the percentage difference from the mean final reconstruction error is shown, illustrating the variability between independent runs.

**Figure S4:**
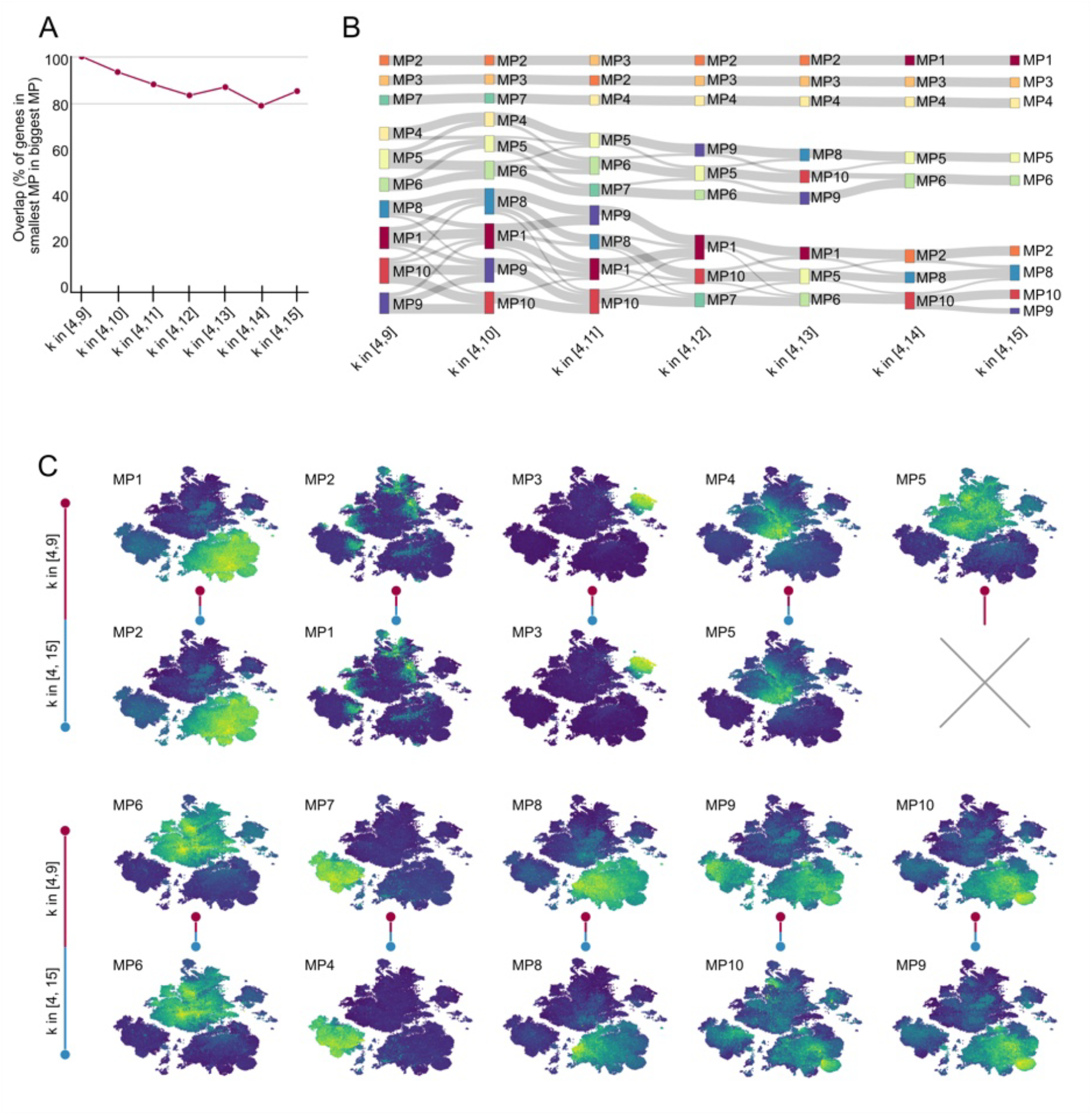
Stability of kaleidoCell meta-programs (MPs) across varying factorization ranges on GBmap. A. Evolution of MP composition across different ranges of *k*. The x-axis shows the range of *k* values used (from [4, 9] to [4, 15]), with 10 MPs defined in each case. MPs are matched across runs using the Hungarian algorithm. Overlap is quantified using the overlap coefficient, defined as the percentage of genes in the smaller MP that are contained in the larger MP. B. Sankey diagram illustrating gene flow between MPs across different k ranges, highlighting the redistribution of genes between matched programs. C. Mapping of MPs between the *k*∈ [4, 9] and *k*∈ [4, 15] configurations, showing correspondences identified via Hungarian matching.

**Figure S5:**
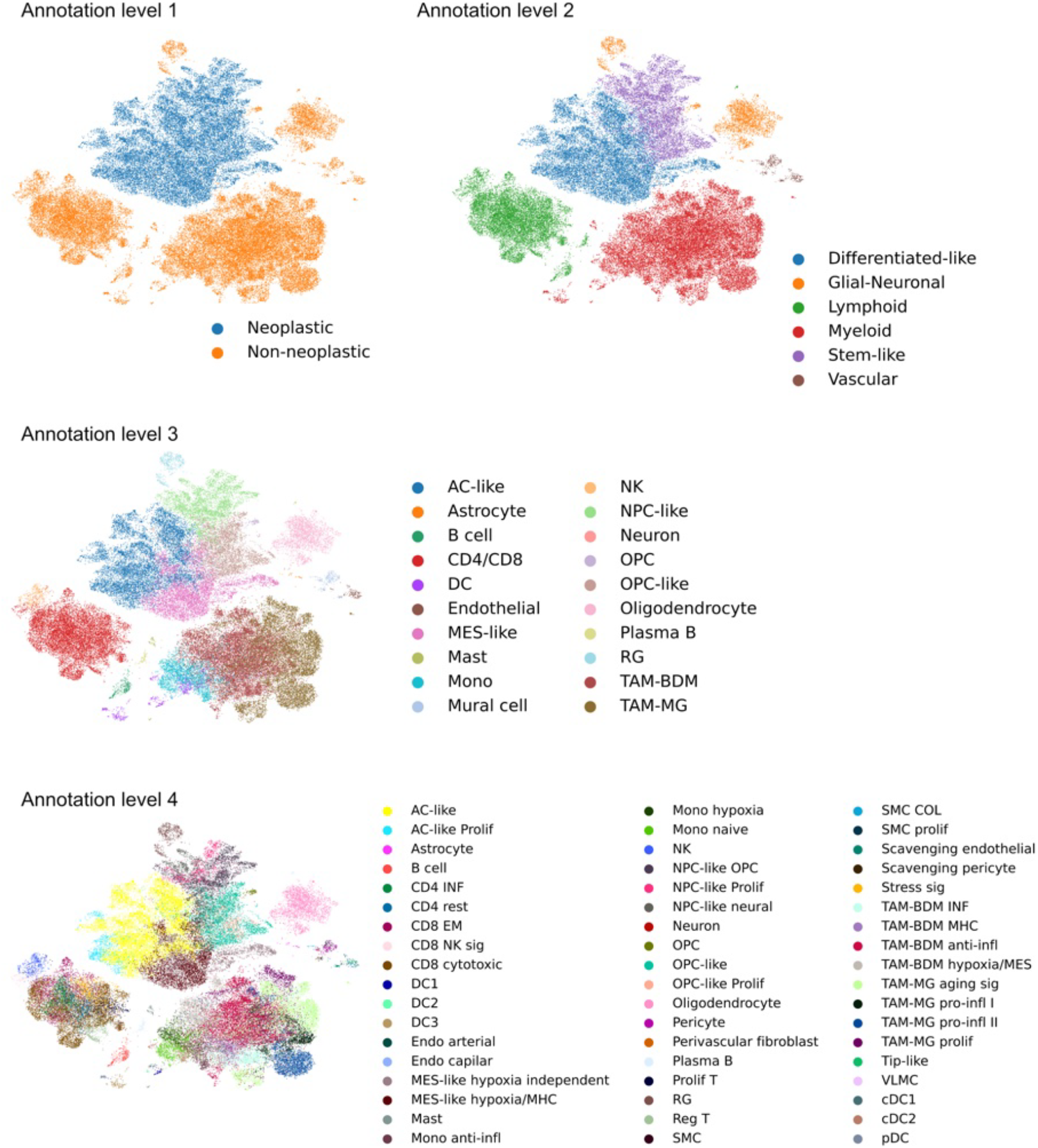
Presentation of the different annotation levels in GBmap.

**Figure S6:**
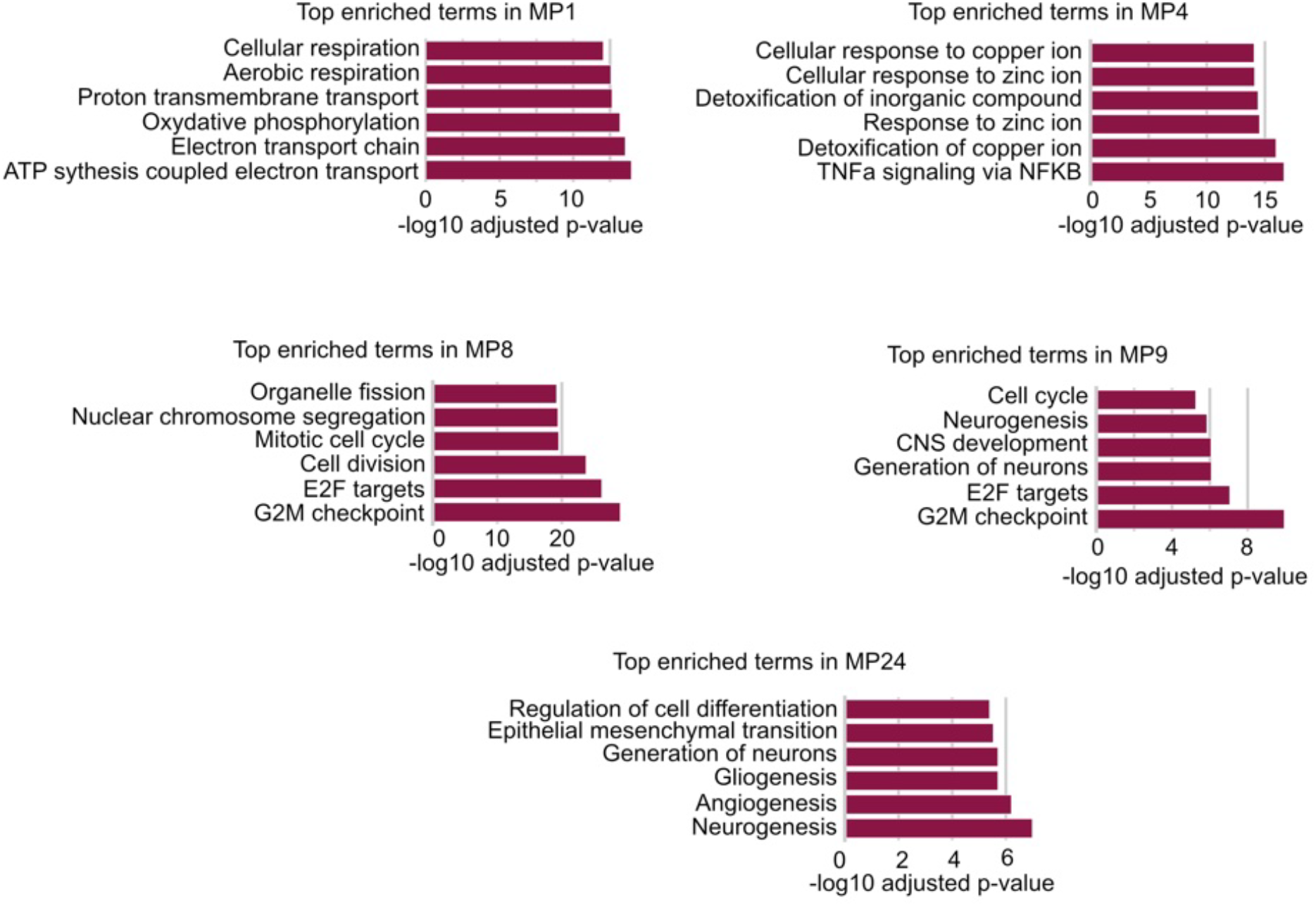
DMSO-Panobinostat meta-programs close up. Gene set enrichment analysis per meta-program (MP).

## References

Kuhn, Harold W (1955). “The Hungarian method for the assignment problem”. In: Naval Research Logistics Quarterly 2.1–2, pp. 83–97. doi: 10.1002/nav.3800020109.

Ward, Joe H (1963). “Hierarchical grouping to optimize an objective function”. In: Journal of the American Statistical Association 58.301, pp. 236–244. doi: 10.1080/01621459.1963.10500845.

Rousseeuw, Peter J (1987). “Silhouettes: a graphical aid to the interpretation and validation of cluster analysis”. In: Journal of Computational and Applied Mathematics 20, pp. 53–65. doi: 10.1016/0377-0427(87)90125-7.

Lee, Daniel D and H Sebastian Seung (1999). “Learning the parts of objects by non-negative matrix factorization”. In: Nature 401.6755, pp. 788–791. doi: 10.1038/44565.

Lee, Daniel D and H Sebastian Seung (2001). “Algorithms for non-negative matrix factorization”. In: Advances in Neural Information Processing Systems. Vol. 13. Neural Information Processing Systems Foundation, pp. 556–562.

Wu, Mark X (2003). “Roles of the stress-induced gene IEX-1 in regulation of cell death and oncogenesis”. In: Apoptosis 8.1, pp. 11–18. doi: 10.1023/A:1021688600370.

Li, Fang et al. (2005). “Myc stimulates nuclearly encoded mitochondrial genes and mitochondrial biogenesis”. In: Molecular and Cellular Biology 25.14, pp. 6225–6234. doi: 10.1128/MCB.25.14.6225-6234.2005.

Kim, Youn Kyung et al. (2006). “Activation of NF-kB by HDAC inhibitor apicidin through Sp1-dependent de novo protein synthesis: its implication for resistance to apoptosis”. In: Cell Death & Differentiation 13.12, pp. 2033–2041. doi: 10.1038/sj.cdd.4401915.

Hunter, John D (2007). “Matplotlib: A 2D Graphics Environment”. In: Computing in Science & Engineering 9.3, pp. 90–95. doi: 10.1109/MCSE.2007.55.

Domingo-Domènech, Joan et al. (2008). “Inactivation of NF-kB by proteasome inhibition contributes to increased apoptosis induced by histone deacetylase inhibitors in human breast cancer cells”. In: Breast Cancer Research and Treatment 112.1, pp. 53–62. doi: 10.1007/s10549-007-9837-8.

McKinney, Wes (2010). “Data Structures for Statistical Computing in Python”. In: Proceedings of the 9th Python in Science Conference. Ed. by Stéfan van der Walt and Jarrod Millman, pp. 51–56. doi: 10.25080/Majora-92bf1922-00a.

Seabold, Skipper and Josef Perktold (2010). “Statsmodels: Econometric and Statistical Modeling with Python”. In: Proceedings of the 9th Python in Science Conference. Ed. by Stéfan van der Walt and Jarrod Millman, pp. 92–96. doi: 10.25080/Majora-92bf1922-011.

Pedregosa, Fabian et al. (2011). “Scikit-learn: Machine Learning in Python”. In: Journal of Machine Learning Research 12, pp. 2825–2830. URL: https://jmlr.org/papers/v12/pedregosa11a.html.

Cornago, Miriam et al. (2014). “Histone deacetylase inhibitors promote glioma cell death by G2 checkpoint abrogation leading to mitotic catastrophe”. In: Cell Death and Disease 5.10, e1435. doi: 10.1038/cddis.2014.412.

Patel, Anoop P et al. (2014). “Single-cell RNA-seq highlights intratumoral heterogeneity in primary glioblastoma”. In: Science 344.6190, pp. 1396–1401. doi: 10.1126/science.1254257.

Marques, Sueli et al. (2016). “Oligodendrocyte heterogeneity in the mouse juvenile and adult central nervous system”. In: Science 352.6291, pp. 1326–1329. doi: 10.1126/science.aaf6463.

Muüller, Sören et al. (2017). “Single-cell profiling of human gliomas reveals macrophage ontogeny as a basis for regional differences in macrophage activation in the tumor microenvironment”. In: Genome Biology 18.1, p. 234. doi: 10.1186/s13059-017-1362-4.

Nebbioso, Angela et al. (2017). “c-Myc Modulation and Acetylation Is a Key HDAC Inhibitor Target in Cancer”. In: Clinical Cancer Research 23.10, pp. 2542–2555. doi: 10.1158/1078-0432.CCR-15-2388.

Hwang, Byungjin, Ji Hyun Lee, and Duhee Bang (2018). “Single-cell RNA sequencing technologies and bioinformatics pipelines”. In: Experimental & Molecular Medicine 50.8, pp. 1–14. doi: 10.1038/s12276-018-0071-8.

Kotliar, Dylan et al. (2019). “Identifying gene expression programs of cell-type identity and cellular activity with single-cell RNA-Seq”. In: eLife 8, e43803. doi: 10.7554/eLife.43803.

Levitin, Hanna Mendes et al. (2019). “De novo gene signature identification from single-cell RNA-seq with hierarchical Poisson factorization”. In: Molecular Systems Biology 15.2, e8557. doi: 10.15252/msb.20188557.

Luecken, Malte D and Fabian J Theis (2019). “Current best practices in single-cell RNA-seq analysis: a tutorial”. In: Molecular Systems Biology 15.6, e8746. doi: 10.15252/msb.20188746.

Neftel, Cyril et al. (2019). “An integrative model of cellular states, plasticity, and genetics for glioblastoma”. In: Cell 178.4, 835–849.e21. doi: 10.1016/j.cell.2019.06.024.

Paszke, Adam et al. (2019). “PyTorch: An imperative style, high-performance deep learning library”. In: Advances in Neural Information Processing Systems 32. Curran Associates, Inc., pp. 8024–8035.

Was, Henryk et al. (2019). “Histone deacetylase inhibitors exert anti-tumor effects on human adherent and stem-like glioma cells”. In: Clinical Epigenetics 11.1, p. 11. doi: 10.1186/s13148-018-0598-5.

Couturier, Charles P et al. (2020). “Single-cell RNA-seq reveals that glioblastoma recapitulates a normal neurodevelopmental hierarchy”. In: Nature Communications 11.1, p. 3406. doi: 10.1038/s41467-020-17186–5.

Harris, Charles R et al. (2020). “Array programming with NumPy”. In: Nature 585.7825, pp. 357–362. doi: 10.1038/s41586-020-2649-2.

Klemm, Florian et al. (2020). “Interrogation of the Microenvironmental Landscape in Brain Tumors Reveals Disease-Specific Alterations of Immune Cells”. In: Cell 181.7, 1643–1660.e17. doi: 10.1016/j.cell.2020.05.007.

Quintero, Andres et al. (2020). “ShinyButchR: Interactive NMF-based decomposition workflow of genome-scale datasets”. In: Biology Methods and Protocols 5.1, bpaa022. doi: 10.1093/biomethods/bpaa022.

Virtanen, Pauli et al. (2020). “SciPy 1.0: fundamental algorithms for scientific computing in Python”. In: Nature Methods 17.3, pp. 261–272. doi: 10.1038/s41592-019-0686-2.

Lou, Q. et al. (2021). “Retinoic Acid Inhibits Tumor-Associated Mesenchymal Stromal Cell Transformation in Melanoma”. In: Frontiers in Cell and Developmental Biology 9, p. 658757. doi: 10.3389/fcell.2021.658757.

Squair, Jordan W et al. (2021). “Confronting false discoveries in single-cell differential expression”. In: Nature Communications 12.1, p. 5692. doi: 10.1038/s41467-021-25960-2.

Waskom, Michael L (2021). “seaborn: statistical data visualization”. In: Journal of Open Source Software 6.60, p. 3021. doi: 10.21105/joss.03021.

Zhao, Wenting et al. (2021). “Deconvolution of cell type-specific drug responses in human tumor tissue with single-cell RNA-seq”. In: Genome Medicine 13.1, p. 82. doi: 10.1186/s13073-021-00894-y.

Barkley, Dalia et al. (2022). “Cancer cell states recur across tumor types and form specific interactions with the tumor microenvironment”. In: Nature Genetics 54.8, pp. 1192–1201. doi: 10.1038/s41588-022-01141-9.

Fang, Zhuoqing, Xinyuan Liu, and Gary Peltz (2023). “GSEApy: a comprehensive package for performing gene set enrichment analysis in Python”. In: Bioinformatics 39.1, btac757. doi: 10.1093/bioinformatics/btac757.

Gavish, Avishai et al. (2023). “Hallmarks of transcriptional intratumour heterogeneity across a thousand tumours”. In: Nature 618.7965, pp. 598–606. doi: 10.1038/s41586-023-06130-4.

Jane, Esther P et al. (2023). “Targeting mitochondrial energetics reverses panobinostat-and marizomib-induced resistance in pediatric and adult high-grade gliomas”. In: Molecular Oncology 17.9, pp. 1821–1843. doi: 10.1002/1878-0261.13427.

Levitin, Hanna Mendes et al. (2023). “Consensus scHPF identifies cell type-specific drug responses in glioma by integrating large-scale scRNA-seq”. In: bioRxiv. doi: 10.1101/2023.12.05.570193.

Takacs, Gabriel P. et al. (2023). “Glioma-derived CCL2 and CCL7 mediate migration of immune suppressive CCR2+/CX3CR1+ M-MDSCs into the tumor microenvironment in a redundant manner”. In: Frontiers in Immunology 13, p. 993444. doi: 10.3389/fimmu.2022.993444.

Virshup, Isaac et al. (2023). “The scverse project provides a computational ecosystem for single-cell omics data analysis”. In: Nature Biotechnology 41.5, pp. 604–606. doi: 10.1038/s41587-023-01733-8.

Kunes, Russell Z et al. (2024). “Supervised discovery of interpretable gene programs from single-cell data”. In: Nature Biotechnology 42.7, pp. 1084–1095. doi: 10.1038/s41587-023-01940-3.

Yerly, Laura et al. (2024). “Wounding triggers invasive progression in human basal cell carcinoma”. In: bioRxiv, pp. 2024–05.

Barsotti, Anne M et al. (2025). “Impaired mitochondrial metabolism is a critical cancer vulnerability for MYC inhibitors”. In: Science Advances 11. doi: 10.1126/sciadv.adw5228.

Cirigliano, S. et al. (2025). “Targeting Glioblastoma Cell State Plasticity for Enhanced Therapeutic Efficacy”. In: bioRxiv. doi: 10.1101/2025.09.08.674897.

Ruiz-Moreno, Cristian et al. (2025). “Charting the single-cell and spatial landscape of IDH-wild-type glioblas-toma with GBmap”. In: Neuro-Oncology 27.9, pp. 2281–2295. doi: 10.1093/neuonc/noaf113.

Song, H., T. Fu, Y. Geng, et al. (2025). “D-Dopachrome Tautomerase-Driven Astrocytic CCL7 Aggravates Neuropathology by Recruitment of Microglia Following Spinal Cord Injury”. In: The FASEB Journal 39.22, e71248. doi: 10.1096/fj.202500902R.

Radig, Jean et al. (2026). “scArchon: a scalable benchmarking framework for assessing single-cell perturbation models”. In: Genome Biology. doi: 10.1186/s13059-026-04104-z.

